# ChAdOx1 nCoV-19 vaccination prevents SARS-CoV-2 pneumonia in rhesus macaques

**DOI:** 10.1101/2020.05.13.093195

**Authors:** Neeltje van Doremalen, Teresa Lambe, Alexandra Spencer, Sandra Belij-Rammerstorfer, Jyothi N. Purushotham, Julia R. Port, Victoria Avanzato, Trenton Bushmaker, Amy Flaxman, Marta Ulaszewska, Friederike Feldmann, Elizabeth R. Allen, Hannah Sharpe, Jonathan Schulz, Myndi Holbrook, Atsushi Okumura, Kimberly Meade-White, Lizzette Pérez-Pérez, Cameron Bissett, Ciaran Gilbride, Brandi N. Williamson, Rebecca Rosenke, Dan Long, Alka Ishwarbhai, Reshma Kailath, Louisa Rose, Susan Morris, Claire Powers, Jamie Lovaglio, Patrick W. Hanley, Dana Scott, Greg Saturday, Emmie de Wit, Sarah C. Gilbert, Vincent J. Munster

## Abstract

Severe acute respiratory syndrome coronavirus-2 (SARS-CoV-2) emerged in December 2019^1,2^ and is responsible for the COVID-19 pandemic^3^. Vaccines are an essential countermeasure urgently needed to control the pandemic^4^. Here, we show that the adenovirus-vectored vaccine ChAdOx1 nCoV-19, encoding the spike protein of SARS-CoV-2, is immunogenic in mice, eliciting a robust humoral and cell-mediated response. This response was not Th2 dominated, as demonstrated by IgG subclass and cytokine expression profiling. A single vaccination with ChAdOx1 nCoV-19 induced a humoral and cellular immune response in rhesus macaques. We observed a significantly reduced viral load in bronchoalveolar lavage fluid and respiratory tract tissue of vaccinated animals challenged with SARS-CoV-2 compared with control animals, and no pneumonia was observed in vaccinated rhesus macaques. Importantly, no evidence of immune-enhanced disease following viral challenge in vaccinated animals was observed. ChAdOx1 nCoV-19 is currently under investigation in a phase I clinical trial. Safety, immunogenicity and efficacy against symptomatic PCR-positive COVID-19 disease will now be assessed in randomised controlled human clinical trials.

---

We previously demonstrated that a single dose of ChAdOx1 MERS, a chimpanzee adeno (ChAd)-vectored vaccine platform encoding the spike protein of MERS-CoV, protected non-human primates (NHPs) against MERS-CoV-induced disease^5^. Here, we designed a ChAdOx1-vectored vaccine encoding a codon-optimised full-length spike protein of SARS-CoV-2 (YP_009724390.1) with a human tPA leader sequence, provisionally named ChAdOx1 nCoV-19, similar to the approach for ChAdOx1 MERS^5^.

## Immunogenicity in mice

Two mouse strains (BALB/c, N=5 and outbred CD1, N=8) were vaccinated intramuscularly (IM) with ChAdOx1 nCoV-19 or ChAdOx1 GFP, a control vaccine expressing green fluorescent protein. Humoral and cellular immunity were studied 9-14 days later. Total IgG titers were detected against spike protein subunits S1 and S2 in all vaccinated mice (Figure 1a). Profiling of the IgG subclasses showed a predominantly Th1 response post vaccination (Extended Data Figure 1a). Virus-specific neutralising antibodies were detected in all mice vaccinated with ChAdOx1 nCoV-19, whereas no neutralisation was detected in serum from mice vaccinated with ChAdOx1 GFP (Figure 1b). Splenic T-cell responses measured by IFN-γ ELISpot and intracellular cytokine staining (ICS) were detected against peptides spanning the full length of the spike construct (Figure 1c). Again, a strong Th1-type response was detected post vaccination as supported by high levels of IFN-γ and TNF-α, and low levels of IL-4 and IL-10 (Figure 1d & Extended Data Figure 1b-c).

**Figure 1:**
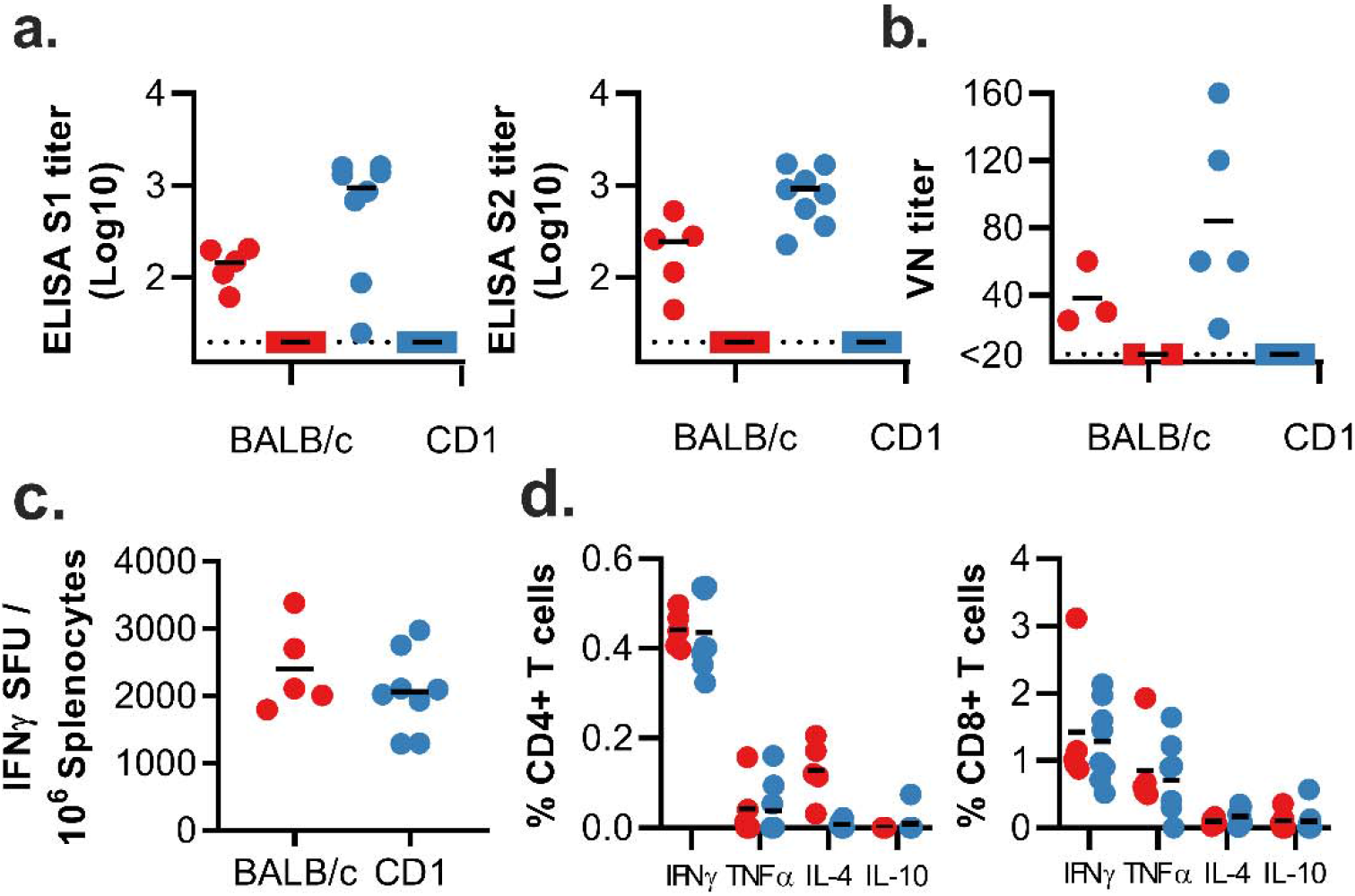
Humoral and cellular immune responses to ChAdOx1 nCoV-19 vaccination in mice. a. End point titer of serum IgG detected against S1 or S2 protein. Control mice were below the limit of detection. b. Virus neutralizing titer in serum. c. Summed IFN-γ ELISpot responses in splenocytes toward peptides spanning the spike protein. Control mice had low (<100 SFU) or no detectable response. d. Summed frequency of spike-specific cytokine positive CD4+ or CD8+ T cells. BALB/c = red; CD1 = blue; vaccinated = circle; control = square; dotted line = limit of detection; line = mean; SFU = spot-forming units.

### Immunogenicity in rhesus macaques

We next evaluated the efficacy of ChAdOx1 nCoV-19 in rhesus macaques, a non-human primate model that displays robust infection of upper and lower respiratory tract and virus shedding upon inoculation with SARS-CoV-2^6^. Twenty-eight days before challenge, 6 animals were vaccinated intramuscularly with 2.5 × 10^10^ ChAdOx1 nCoV-19 virus particles each, half of the dose currently administered in human clinical trials. As a control, three animals were vaccinated via the same route with the same dose of ChAdOx1 GFP (Figure 2a). Spike-specific antibodies were present as early as 14 days post vaccination and endpoint IgG titers of 400-6400 were measured on the day of challenge (Figure 2b). Virus-specific neutralising antibodies were detectable in all vaccinated animals before challenge (VN titer = 5-40), whereas no virus-specific neutralising antibodies were detected in control animals (Figure 2c). Finally, SARS-CoV-2 spike specific T-cell responses were detected by IFN-γ ELISpot assay and involved stimulation of peripheral blood mononuclear cells (PBMCs) with a peptide library spanning the full length of the spike protein (Figure 2d).

**Figure 2.**
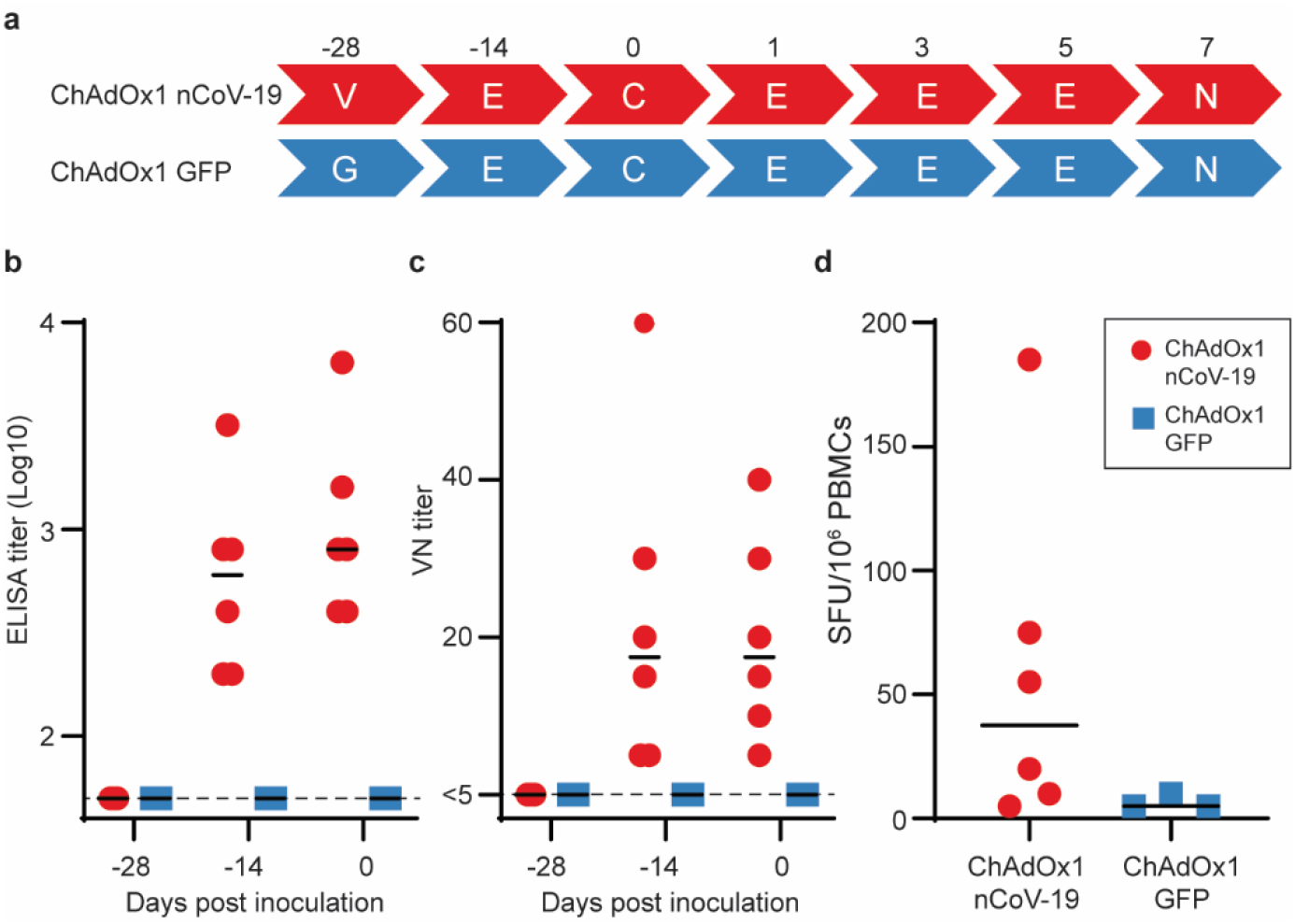
Humoral and cellular immune responses to ChAdOx1 nCoV-19 vaccination in rhesus macaques. a. Study schedule for NHPs. V = vaccination with ChAdOx1 nCoV-19; G = vaccination with ChAdOx1 GFP; E = exam; N = exam and necropsy. b. Virus neutralizing titer in serum. d. Summed IFN-γ ELISpot responses in PBMCs toward peptides spanning the spike protein Vaccinated animals = red circles; control animals = blue squares; dotted line = limit of detection; line – median; SFU = spot-forming units.

### Clinical signs

Upon challenge with 2.6 × 10^6^ TCID_50_ SARS-CoV-2 to both the upper and lower respiratory tract the average clinical score of control animals was higher compared to ChAdOx1 nCoV-19 vaccinated animals. This was significantly different as determined via Mann-Whitney’s rank’s test on 3 and 5 DPI (Figure 3a). All control animals showed an increase in respiratory rate, compared to 3 out of 6 vaccinated animals. Respiratory signs persisted longer in control animals (Extended Data Table 1).

**Figure 3.**
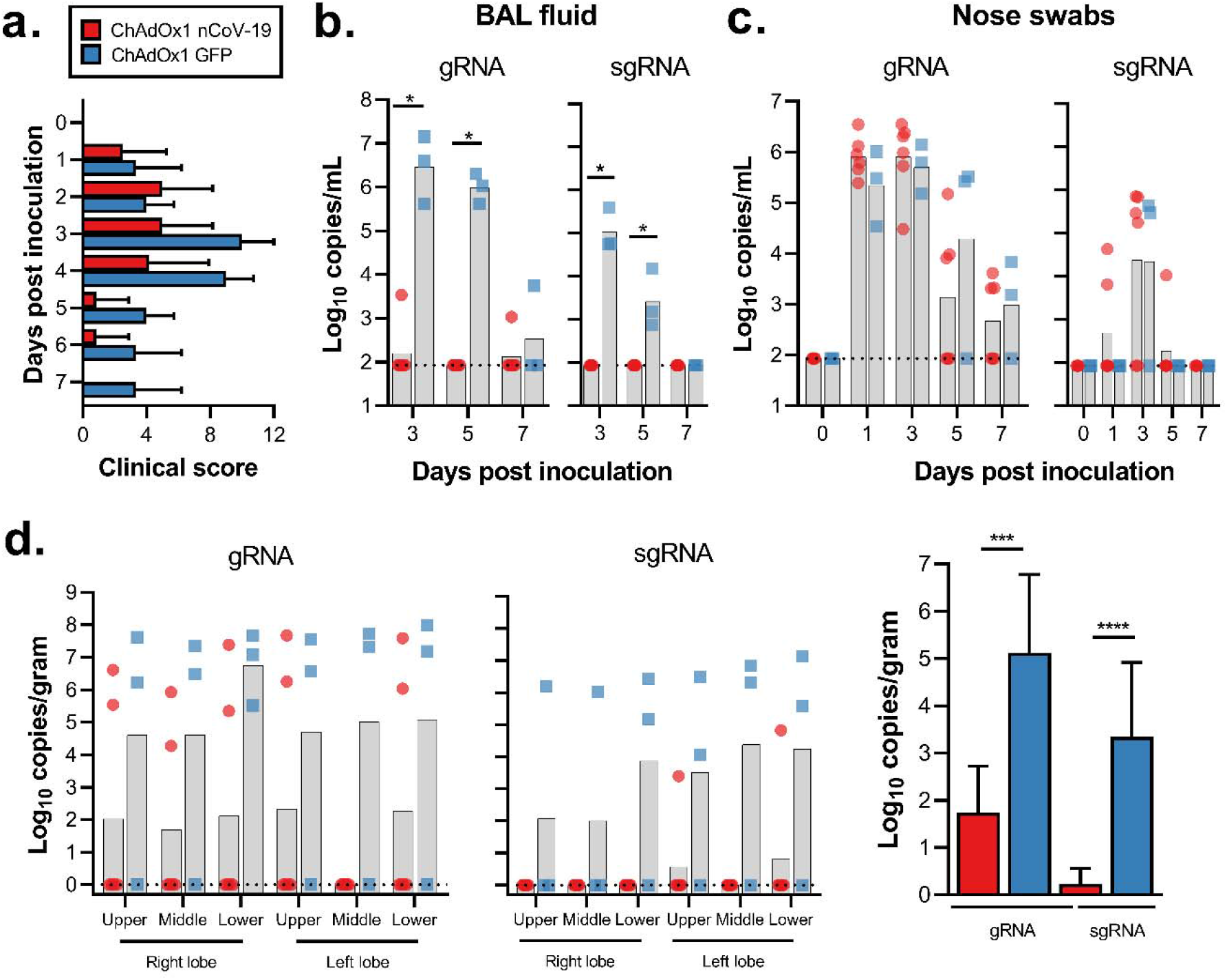
Clinical signs and viral load in rhesus macaques inoculated with SARS-CoV-2 after vaccination with ChAdOx1 nCoV-19. a. Mean clinical score with standard deviation in NHPs. Any scoring associated with food was removed from final score. b. Viral load in BAL fluid obtained from rhesus macaques, bar at geometric mean. *=p-value<0.0166. c. Viral load in nose swabs obtained from rhesus macaques, bar at geometric mean. d. Viral load in tissues at 7 DPI. Pictured are individual values with geometric mean bars (left panels) and geometric mean of all lung lobes per group (right panel). ***=p-value<0.001; ****=p-value<0.0001. Vaccinated animals = red circles; control animals = blue squares; dotted line = limit of detection.

### Viral load in respiratory tract samples

In the control animals, viral genomic RNA (gRNA) was detected in BAL fluid on all days and viral subgenomic RNA (sgRNA), indicative of virus replication, was detected at 3 and 5 days post inoculation (DPI). In contrast, viral gRNA was detected in only two animals, either on 3 or 7 DPI, and no viral sgRNA could be detected in BAL fluid obtained from vaccinated animals (p=0.0119, Figure 3b). Viral gRNA was detected in nose swabs from all animals and no difference in viral load in nose swabs was found on any days between vaccinated and control animals (Figure 3c).

### Cytokine response

Cytokines in serum were analysed after challenge to monitor immune responses. We observed an upregulation in IFN-γ at 1 DPI in ChAdOx1 nCoV-19 vaccinated animals, but not in control animals. No significant differences were observed between ChAdOx1 nCoV-19 and control animals for TNF-α, IL-2, IL-4, IL-6, and IL-10 (Extended Data Figure 2).

### Pulmonary pathology

At 7 days post inoculation, all animals were euthanized, and tissues were collected. None of the vaccinated monkeys developed pulmonary pathology after inoculation with SARS-CoV-2. All lungs were histologically normal and no evidence of viral pneumonia nor immune-enhanced inflammatory disease was observed. In addition, no SARS-CoV-2 antigen was detected by immunohistochemistry in the lungs of any of the vaccinated animals. Two out of 3 control animals developed some degree of viral interstitial pneumonia. Lesions were widely separated and characterized by thickening of alveolar septae by small amounts of edema fluid and few macrophages and lymphocytes. Alveoli contained small numbers of pulmonary macrophages and, rarely, edema. Type II pneumocyte hyperplasia was observed. Multifocally, perivascular infiltrates of small numbers of lymphocytes forming perivascular cuffs were observed. Immunohistochemistry demonstrated viral antigen in type I and II pneumocytes, as well as in alveolar macrophages (Figure 4).

**Figure 4.**
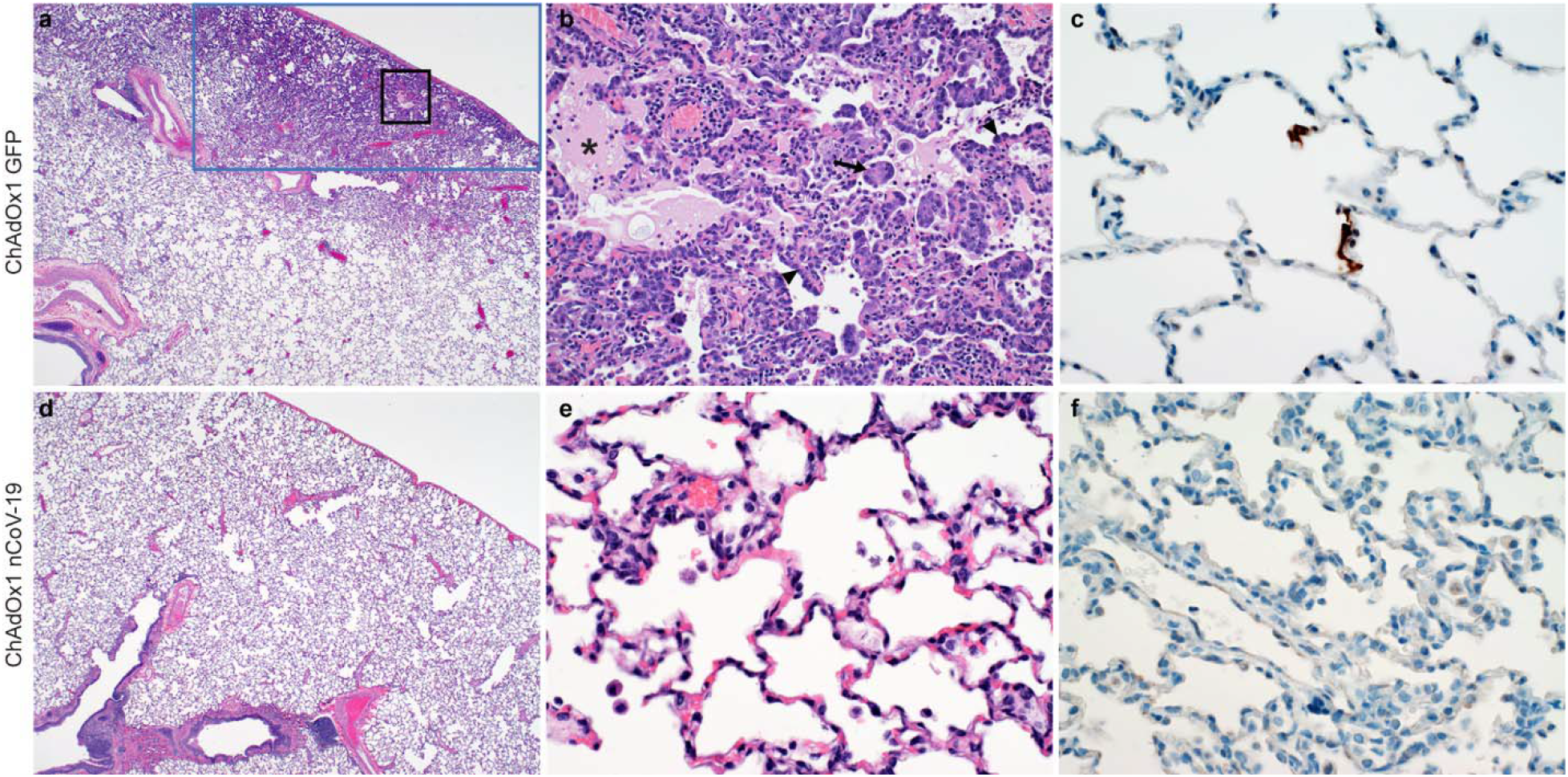
Histological changes in lungs of rhesus macaques on 7 dpi. a) Focal interstitial pneumonia in lungs of a control animal (blue box). The area in the black box is magnified in panel b. b) Interstitial pneumonia with edema (asterisk), type II pneumocyte hyperplasia (arrowhead) and syncytial cells (arrow) in control animals. c) SARS-CoV-2 antigen (visible as red-brown staining) was detected by immunohistochemistry in type I and type II pneumocytes in the lungs of control animals. d) No histological changes were observed in the lungs of ChadOx1 nCoV-19-vaccinated animals. e) Higher magnification of lung tissue in panel d. No evidence of pneumonia or immune-enhanced inflammation is observed. f) No SARS-CoV-2 antigen was detected by immunohistochemistry in the lungs of vaccinated animals. Magnification: panels a, d 40x; panels b, c, e, f 400x.

### Viral load in respiratory tract

Viral gRNA load was high in lung tissue of control animals and viral sgRNA was detected in 2 out of 3 control animals (Figure 3d). In contrast, the viral gRNA load was significantly lower in lung tissue obtained from vaccinated animals as determined via Mann-Whitney’s rank test and below limits of detection in two vaccinated animals. Viral sgRNA was detected in lung tissue obtained from 1 out of 6 vaccinated animals (p<0.0001, Figure 3d). Viral gRNA could be detected in other tissues but was low in both groups (Extended Data Figure 3).

## Discussion

Here, we showed that a single vaccination with ChAdOx1 nCoV-19 is effective in preventing damage to the lungs upon high dose challenge with SARS-CoV-2. Similarly a recent study showed that a triple vaccination regime of a high dose of whole inactivated SARS-CoV-2 protected rhesus macaques from SARS-CoV-2 pneumonia^7^.

Viral loads in BAL fluid and lung tissue of vaccinated animals were significantly reduced, suggesting that vaccination prevents virus replication in the lower respiratory tract. Despite this marked difference in virus replication in the lungs, reduction in viral shedding from the nose was not observed. However, animals were challenged with a high dose of virus via multiple routes, which likely does not reflect a realistic human exposure. Whether a lower challenge dose would result in more efficient protection of the upper respiratory tract remains to be determined.

Several preclinical studies of vaccines against SARS-CoV-1 resulted in immunopathology after vaccination and challenge, with more severe disease in vaccinated animals than in controls^8–10^. Importantly, we did not see any evidence of immune-enhanced disease in vaccinated animals. The immune response was not skewed towards a Th2 response in mice nor in NHPs, there was no increase in clinical signs or virus replication throughout the study in vaccinated NHPs compared to controls and no markers of disease enhancement in lung tissue of NHPs, such as an influx of neutrophils were observed. These data informed the start of the phase I clinical trial with ChAdOx1 nCoV-19 on April 23, 2020. As of May 13, 2020, more than 1000 volunteers have participated in the clinical trials. This study is thus an important step towards the development of a safe and efficacious SARS-CoV-2 vaccine.

## Methods

### Ethics Statement

Mice – Mice were used in accordance with the UK Animals (Scientific Procedures) Act under project license number P9804B4F1 granted by the UK Home Office. Animals were group housed in IVCs under SPF conditions, with constant temperature and humidity with lighting on a fixed light/dark cycle (12-hours/12-hours).

NHPs – Animal experiment approval was provided by the Institutional Animal Care and Use Committee (IACUC) at Rocky Mountain Laboratories. Animal experiments were executed in an Association for Assessment and Accreditation of Laboratory Animal Care (AALAC)-approved facility by certified staff, following the basic principles and guidelines in the NIH Guide for the Care and Use of Laboratory Animals, the Animal Welfare Act, United States Department of Agriculture and the United States Public Health Service Policy on Humane Care and Use of Laboratory Animals. Rhesus macaques were housed in individual primate cages allowing social interactions, in a climate-controlled room with a fixed light/dark cycle (12-hours/12-hours) and monitored a minimum of twice daily. Commercial monkey chow, treats, and fruit were provided by trained personnel. Water was available ad libitum. Environmental enrichment consisted of a variety of human interaction, commercial toys, videos, and music. The Institutional Biosafety Committee (IBC) approved work with infectious SARS-CoV-2 virus strains under BSL3 conditions. All sample inactivation was performed according to IBC approved standard operating procedures for removal of specimens from high containment.

### Generation of vaccine ChAdOx1 nCoV-19

The spike protein of SARS-CoV-2 (Genbank accession number YP_009724390.1), the surface glycoprotein responsible for receptor binding and fusion/entry into the host cell, was codon optimised for expression in human cell lines and synthesised with the tissue plasminogen activator (tPA) leader sequence at the 5’ end by GeneArt Gene Synthesis (Thermo Fisher Scientific). The sequence, encoding SARS-CoV-2 amino acids 2-1273 and tPA leader, was cloned into a shuttle plasmid using InFusion cloning (Clontech). The shuttle plasmid encodes a modified human cytomegalovirus major immediate early promoter (IE CMV) with tetracycline operator (TetO) sites, poly adenylation signal from bovine growth hormone (BGH), between Gateway® recombination cloning sites. ChAdOx1 nCoV-19 was prepared using Gateway® recombination technology (Thermo Fisher Scientific) between the shuttle plasmid described and the ChAdOx1 destination DNA BAC vector described in^11^ resulting in the insertion of the SARS-CoV-2 expression cassette at the E1 locus. The ChAdOx1 adenovirus genome was excised from the BAC using unique PmeI sites flanking the adenovirus genome sequence. The virus was rescued and propagated in T-Rex 293 HEK cells (Invitrogen) which repress antigen expression during virus propagation. Purification was by CsCl gradient ultracentrifugation. Virus titers were determined by hexon immunostaining assay and viral particles calculated based on spectrophotometry^12,13^.

### Study design animal experiments

Mice – Female BALB/cOlaHsd (BALB/c) (Envigo) and outbred Crl:CD1(ICR) (CD1) (Charles River) mice of at least 6 weeks of age, were immunized IM in the musculus tibialis with 6×10^9^ VP of ChAdOx1 nCoV-19 unless otherwise stated.

NHPs – 9 adult rhesus macaques (8M, 1F) were randomly divided into two groups of six and three animals. Group 1 was vaccinated with ChAdOx1 nCoV-19 at −28 DPI, group 2 was vaccinated with ChAdOx1 GFP at −28 DPI. All vaccinations were done with 2.5 × 10^10^ VP/animal diluted in sterile PBS. Blood samples were obtained before vaccination and 14 days thereafter. Animals were challenged with SARS-CoV-2 strain nCoV-WA1-2020 (MN985325.1) diluted in sterile DMEM on 0 DPI; with administration of 4 mL intratracheally, 1 mL intranasally, 1 mL orally and 0.5 mL ocularly of 4 × 10^5^ TCID_50_/mL virus suspension. Animals were scored daily by the same person who was blinded to study group allocations using a standardized scoring sheet. Clinical exams were performed on −28, −14, 0, 1, 3, and 5 and 7 DPI. Nasal swabs and blood were collected at all exam dates. BAL was performed on 3, 5, and 7 DPI by insertion of an endotracheal tube and bronchoscope into the trachea, then past the 3^rd^ bifurcation, and subsequent installation of 10 mL of sterile saline. Manual suction was applied to retrieve the BAL sample. Necropsy was performed on 7 DPI and the following tissues were collected: cervical lymph node, mediastinal lymph node, conjunctiva, nasal mucosa, oropharynx, tonsil, trachea, all six lung lobes, right and left bronchus, heart, liver, spleen, kidney, stomach, duodenum, jejunum, ileum, cecum, colon, urinary bladder.

### Cells and virus

SARS-CoV-2 strain nCoV-WA1-2020 (MN985325.1) was provided by CDC, Atlanta, USA. Virus propagation was performed in VeroE6 cells in DMEM supplemented with 2% fetal bovine serum, 1 mM L-glutamine, 50 U/ml penicillin and 50 μg/ml streptomycin. VeroE6 cells were maintained in DMEM supplemented with 10% fetal bovine serum, 1 mM L-glutamine, 50 U/ml penicillin and 50 μg/ml streptomycin.

### Virus neutralization assay

Sera were heat-inactivated (30 min, 56 °C), two-fold serial dilutions were prepared in 2% DMEM and 100 TCID_50_ of SARS-CoV-2 was added. After 1hr incubation at 37 °C and 5% CO2, virus:serum mixture was added to VeroE6 cells and incubated at 37°C and 5% CO2. At 5 dpi, cytopathic effect was scored. The virus neutralization titer was expressed as the reciprocal value of the highest dilution of the serum which still inhibited virus replication.

### RNA extraction and quantitative reverse-transcription polymerase chain reaction

Tissues (up to 30 mg) were homogenized in RLT buffer and RNA was extracted using the RNeasy kit (Qiagen) according to the manufacturer's instructions. RNA was extracted from BAL fluid and nasal swabs using the QiaAmp Viral RNA kit (Qiagen) according to the manufacturer's instructions. Viral gRNA^14^ and sgRNA^15^ specific assays were used for the detection of viral RNA. Five μl RNA was tested with the Rotor-GeneTM probe kit (Qiagen) according to instructions of the manufacturer. Dilutions of SARS-CoV-2 standards with known genome copies were run in parallel.

### Enzyme-linked immunosorbent assay for mouse sera

MaxiSorp plates (Nunc) were coated with S1 or S2 (The Native Antigen Company; 50 ng/well) in PBS for overnight adsorption at 4°C. Plates were washed in PBS/Tween (0.05% v/v) and wells blocked using casein (ThermoFisher Scientific) for 1hr at RT. Serially diluted mouse serum samples were added and incubated overnight at 4°C. Plates were washed and Alkaline Phosphatase-conjugated goat anti-mouse IgG (Sigma) was added to all wells for 1hr at RT. After washing pNPP substrate (Sigma) was added. Optical density (OD) values for each well were measured at 405 nm. Endpoint titers were calculated as follows: the log_10_ OD against log_10_ sample dilution was plotted and a regression analysis of the linear part of this curve allowed calculation of the endpoint titer with an OD of three times the background. The same calculation was used for diluting the sera to the same amounts of total IgG for further testing on different IgG subclasses with anti-mouse IgG subclass-specific antibodies (Abcam). The results of the IgG subclass ELISA are presented using OD values.

### Enzyme-linked immunosorbent assay for NHP sera

Stabilized SARS-CoV-2 spike protein was obtained from the Vaccine Research Centre, Bethesda, USA. Maxisorp plates (Nunc) were coated overnight at 4°C with 100 ng/well spike protein in PBS. Plates were blocked with 100 μl of casein in PBS (Thermo Fisher) for 1hr at RT. Serum serially diluted 2x in casein in PBS was incubated at RT for 1hr. Antibodies were detected using affinity-purified polyclonal antibody peroxidase-labeled goat-anti-monkey IgG (Seracare, 074-11-021) in casein and TMB 2-component peroxidase substrate (Seracare, 5120-0047), developed for 5-10 min, and reaction was stopped using stop solution (Seracare, 5150-0021) and read at 450 nm. All wells were washed 4x with PBST 0.1% tween in between steps. Threshold for positivity was set at 3x OD value of negative control (serum obtained from non-human primates prior to start of the experiment) or 0.2, whichever one was higher.

### ELISpot assay and ICS analysis

Single cell suspension of murine splenocytes were prepared by passing cells through 70μM cell strainers and ACK lysis prior to resuspension in complete media. Rhesus macaque PBMCs were isolated from ethylene diamine tetraaceticacid (EDTA) whole blood using LeucosepTM tubes (Greiner Bio-one International GmbH) and Histopaque®-1077 density gradient cell separation medium (Sigma-Aldrich) according to the manufacturers’ instructions.

Mice – For analysis of IFN-γ production by ELISpot, cells were stimulated with pools of S1 or S2 peptides (final concentration of 2μg/ml) on IPVH-membrane plates (Millipore) coated with 5μg/ml anti-mouse IFN-γ (AN18). After 18-20 hours of stimulation, IFN-γ spot forming cells (SFC) were detected by staining membranes with anti-mouse IFN-γ biotin (1μg/ml) (R46A2) followed by streptavidin-Alkaline Phosphatase (1μg/ml) and development with AP conjugate substrate kit (BioRad, UK).

For analysis of intracellular cytokine production, cells were stimulated at 37°C for 6 hours with 2μg/ml S1 or S2 pools of peptide, media or cell stimulation cocktail (containing PMA-Ionomycin, Biolegend), together with 1μg/ml Golgi-plug (BD) with the addition of 2μl/ml CD107a-Alexa647. Cell supernatant was collected and frozen at –20::J C for subsequent analysis by MesoScaleDiscovery (MSD) assay (see below). Following surface staining with CD4-BUV496, CD8-PerCPCy5.5, CD62L-BV711 and CD127-BV650, cells were fixed with 4% paraformaldehyde and stained intracellularly with TNF-α-A488, IL-2-PECy7, IL-4-BV605, IL-10-PE and IFN-γ-e450 diluted in Perm-Wash buffer (BD).

Sample acquisition was performed on a Fortessa (BD) and data analyzed in FlowJo v9 or FlowJo V10 (TreeStar). An acquisition threshold was set at a minimum of 5000 events in the live CD3^+^ gate. Antigen specific T cells were identified by gating on LIVE/DEAD negative, doublet negative (FSC-H vs FSC-A), size (FSC-H vs SSC), CD3^+^, CD4^+^ or CD8^+^ cells and cytokine positive. Cytokine positive responses are presented after subtraction of the background response detected in the corresponding unstimulated sample (media containing CD107a and Golgi-plug) of each individual spleen sample.

NHPs – IFN-γ ELISpot assay of PBMCs was performed using the ImmunoSpot® Human IFN- γ Single-Color Enzymatic ELISpot Assay Kit according to the manufacturer’s protocol (Cellular Technology Limited). PBMCs were plated at a concentration of 100,000 cells per well and were stimulated with four contiguous peptide pools spanning the length of the SARS-CoV-2 spike protein sequence at a concentration of 2 μg/mL per peptide (Mimotopes). ELISpot plates were subjected to overnight formalin inactivation prior to removal from BSL4 for reading. Analysis was performed using the CTL ImmunoSpot® Analyzer and ImmunoSpot® Software (Cellular Technology Limited). Spot forming units (SFU) per 1.0×10^6^ PBMCs were summed across the 4 peptide pools for each animal.

### Measurement of cytokines and chemokines

Mouse samples were assayed using MSD Technology V-PLEX Mouse Cytokine 29-Plex kit according to the manufacturer’s instructions. Non-human primate samples were inactivated with γ-radiation (2 MRad) according to standard operating procedures and assayed on a Bio-Plex 200 instrument (Bio-Rad) using the Non-Human Primate Cytokine MILLIPLEX map 23-plex kit (Millipore) according to the manufacturer’s instructions. LLOD was used for all undetectable and extrapolated values. Only data for cytokines consistently above the lower limit of quantification were included in further anlayses.

Log_10_ Fold Change (Log_10_FC) for mouse samples was calculated as follows:

Log_10_FC = Log_10_((Stimulated (pg/ml) + 1) / (Unstimulated (pg/ml) + 1))

Fold change for NHP samples was calculated as follows:

FC = Concentration (pg/mL) on DX (1, 3, 5, or 7)/Concentration (pg/mL) on D0

### Histology and immunohistochemistry

Necropsies and tissue sampling were performed according to IBC-approved protocols. Lungs were perfused with 10% formalin and processed for histologic review. Harvested tissues were fixed for eight days in 10% neutral-buffered formalin, embedded in paraffin, processed using a VIP-6 Tissue Tek (Sakura Finetek, USA) tissue processor, and embedded in Ultraffin paraffin polymer (Cancer Diagnostics, Durham, NC). Samples were sectioned at 5 μm, and resulting slides were stained with hematoxylin and eosin. Specific anti-CoV immunoreactivity was detected using an in-house SARS-CoV-2 nucleocapsid protein rabbit antibody (Genscript) at a 1:1000 dilution. The IHC assay was carried out on a Discovery ULTRA automated staining instrument (Roche Tissue Diagnostics) with a Discovery ChromoMap DAB (Ventana Medical Systems) kit. All tissue slides were evaluated by a board-certified veterinary anatomic pathologist blinded to study group allocations.

### Statistical analyses

Two-tailed Mann-Whitney’s rank tests were conducted to compare differences between groups. A Bonferroni correction was used to control for type I error rate where required.

### Data availability

Data have been deposited in Figshare: 10.6084/m9.figshare.12290696

## Acknowledgements

The authors would like to acknowledge Olubukola Abiona, Brandon Bailes, Aaron Carmody, Kizzmekia Corbett, Kathleen Cordova, Jayne Faris, Heinz Feldmann, Susan Gerber, Barney Graham, Elaine Haddock, Ryan Kissinger, Michael Jones, Mary Marsh, Kay Menk, Anita Mora, Stephanie Seifert, Les Shupert, Brian Smith, Natalie Thornburg, Amanda Weidow, Marissa Woods, and Kwe Claude Yinda for their contributions to this study. This work was supported by the Intramural Research Program of the National Institute of Allergy and Infectious Diseases (NIAID), National Institutes of Health (NIH) (1ZIAAI001179-01) and the Department of Health and Social Care using UK Aid funding managed by the NIHR.

## Author Information

### Author contributions

NvD, TL, SG and VJM designed the study; NvD, TL, AS, SBR, JNP, JRP, VA, TB, AF, MU, FF, EA, HS, JS, MH, AO, KMW, LPP, CB, CG, BNW, RR, DL, AI, RK, LR, SM, CP, JL, PH, DS, GS, EdW, SG, VJM acquired, analysed and interpreted the data; NvD, TL, EdW, SG, and VJM wrote the manuscript. All authors have approved the submitted version.

## Competing interests

SCG is a board member of Vaccitech and named as an inventor on a patent covering use of ChAdOx1-vectored vaccines and a patent application covering a SARS-CoV-2 (nCoV-19) vaccine. Teresa Lambe is named as an inventor on a patent application covering a SARS-CoV-2 (nCoV-19) vaccine. The remaining authors declare no competing interests.

## Extended Data Legends

**Extended Data Figure 1.**
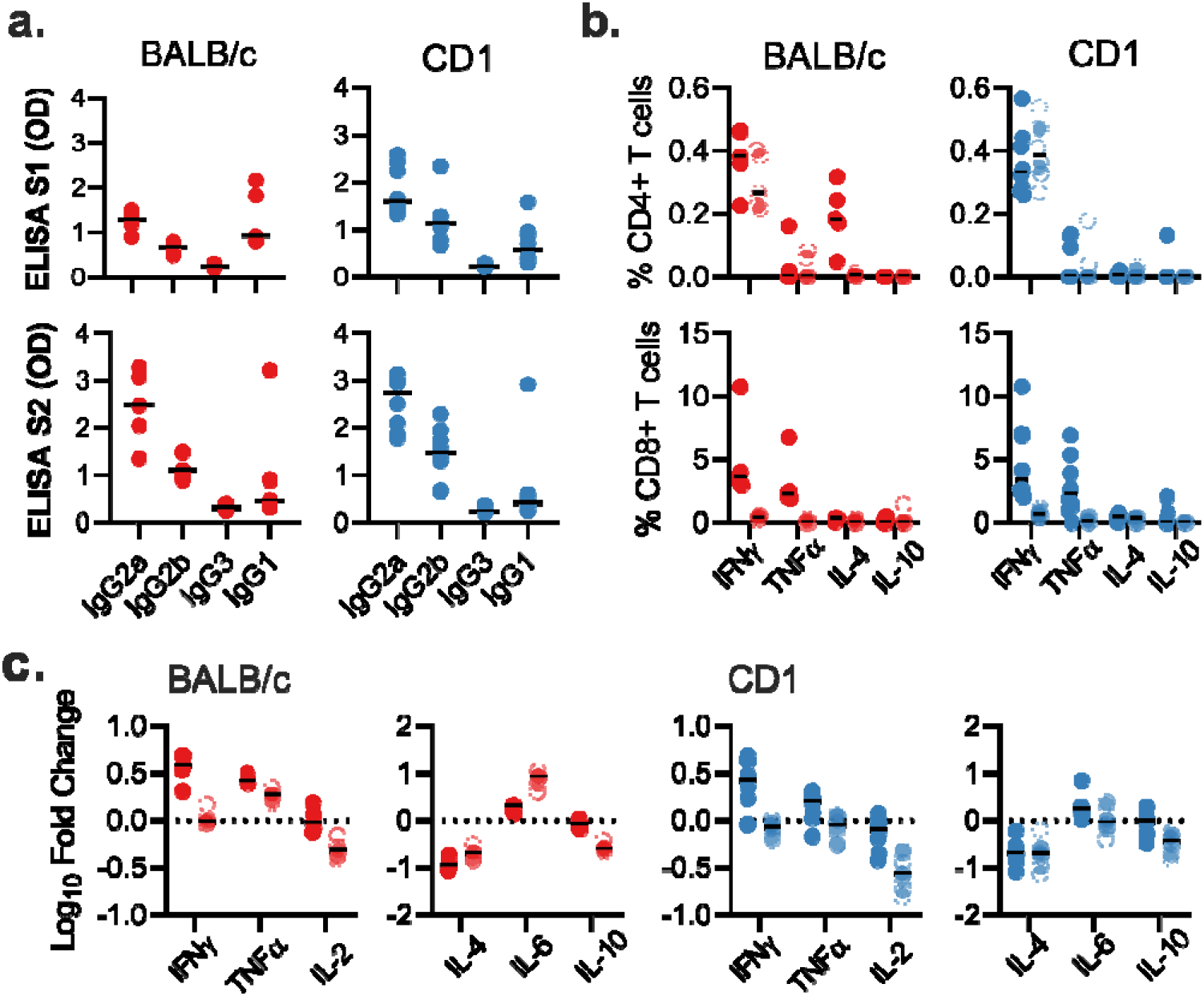
Antigen specific responses following ChAdOx1 nCov19 vaccination. a. IgG subclass antibodies detected against S1 or S2 protein in sera of BALB/c or CD1 mice. b. Frequency of cytokine positive CD4+ or CD8+ T cells following stimulation of splenocytes with S1 pool (dark) or S2 pool (transparent) peptides in BALB/c (red) and CD1 (blue) mice. d. Log10 fold change in cytokine levels in supernatant from S1 (dark) and S2 (transparent) stimulated splenocytes when compared to corresponding unstimulated splenocyte sample for BALB/c and CD1 mice.

**Extended Data Table 1.**
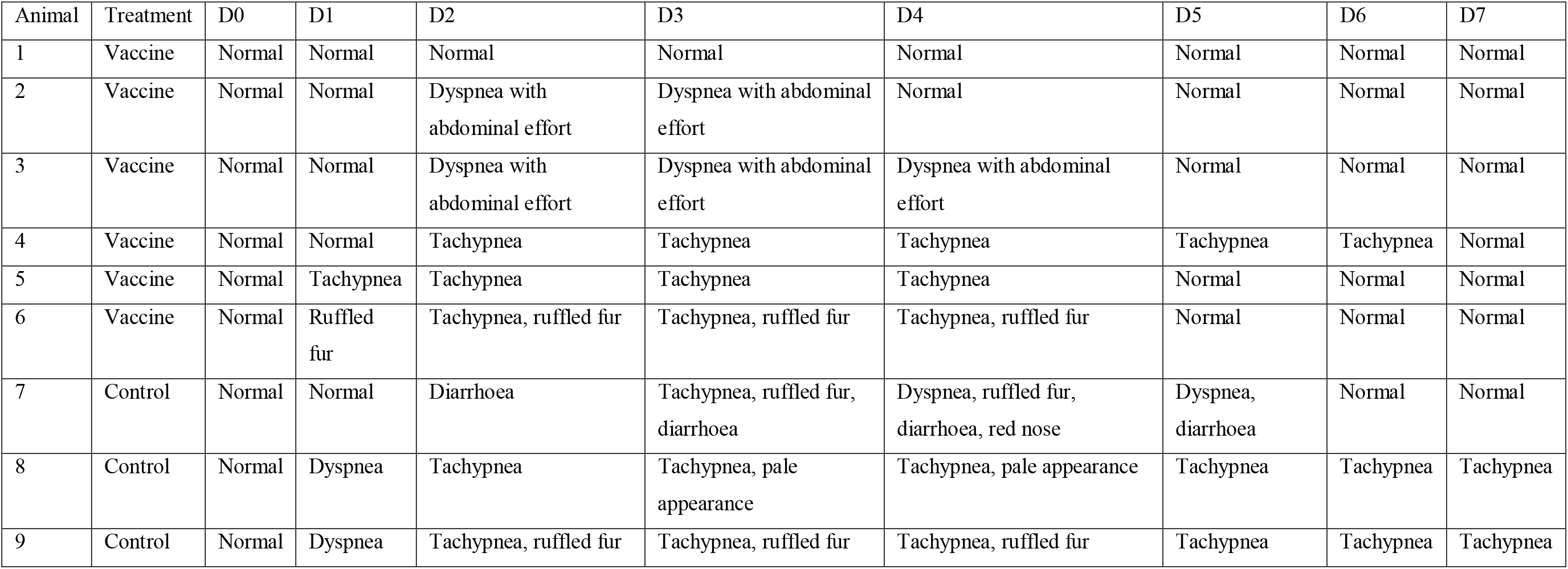
Clinical signs observed in rhesus macaques inoculated with SARS-CoV-2.

**Extended Data Figure 2.**
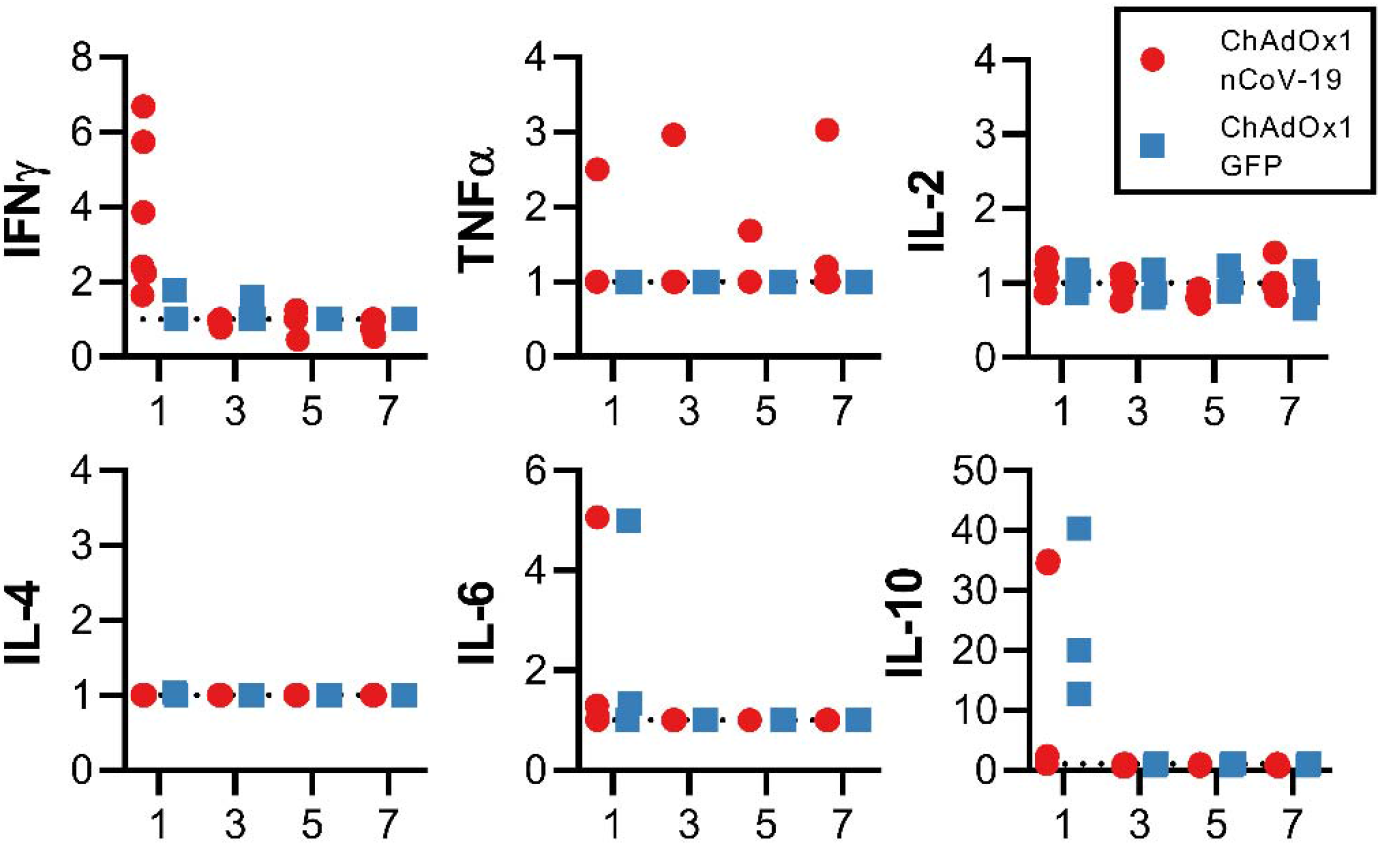
Serum cytokines in rhesus macaques challenged with SARS-CoV-2. Fold increase in cytokines in serum compared to 0 DPI values.

**Extended Data Figure 3.**
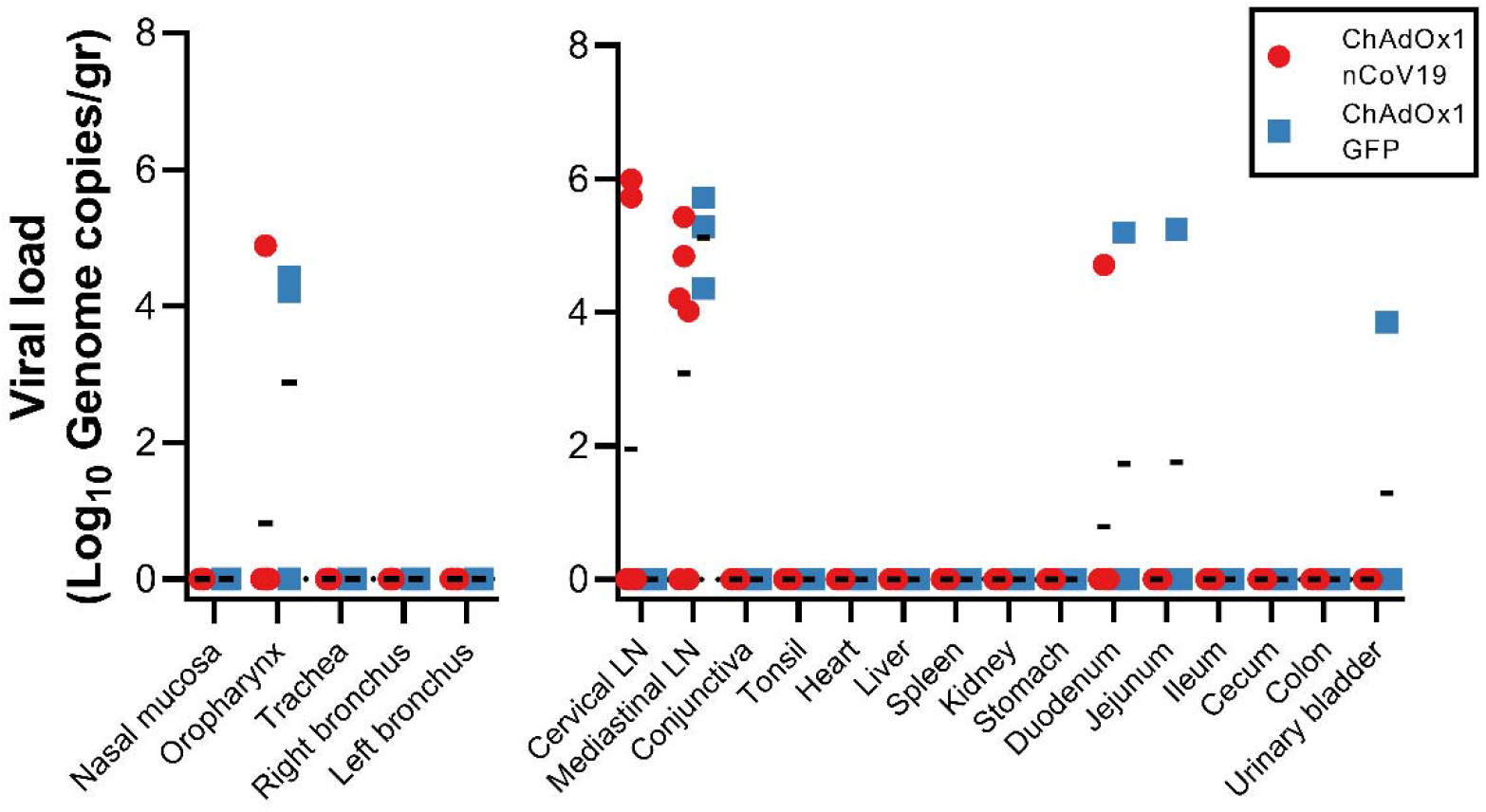
Viral load in rhesus macaques challenged with SARS-CoV-2. Viral genomic RNA in respiratory tissues excluding lung tissue (left panel) and other tissues (right panel). A two-tailed Mann-Whitney’s rank test was performed to investigate statistical significance. Bonferroni correction was applied, and thus statistical significance was reached at p>0.0125.

## References

1 Zhu, N. et al. A Novel Coronavirus from Patients with Pneumonia in China, 2019. N Engl J Med 382, 727–733, doi:10.1056/NEJMoa2001017 (2020).

2 Wu, F. et al. A new coronavirus associated with human respiratory disease in China. Nature 579, 265–269, doi:10.1038/s41586-020-2008-3 (2020).

3 WHO. Coronavirus disease (COVID-19) Situation Report 113, <https://www.who.int/docs/default-source/coronaviruse/situation-reports/20200512-covid-19-sitrep-113.pdf?sfvrsn=feac3b6d_2> (2020).

4 Lurie, N., Saville, M., Hatchett, R. & Halton, J. Developing Covid-19 Vaccines at Pandemic Speed. N Engl J Med, doi:10.1056/NEJMp2005630 (2020).

5 van Doremalen, N. et al. A single dose of ChAdOx1 MERS provides protective immunity in rhesus macaques. Science Advances, doi:10.1126/sciadv.aba8399 (2020).

6 Munster, V. J. et al. Respiratory disease in rhesus macaques inoculated with SARS-CoV-2. Nature, doi:10.1038/s41586-020-2324-7 (2020).

7 Qiang, G. et al. Development of an inactivated vaccine candidate for SARS-CoV-2. Science, doi:10.1126/science.abc1932 (2020).

8 Weingartl, H. et al. Immunization with modified vaccinia virus Ankara-based recombinant vaccine against severe acute respiratory syndrome is associated with enhanced hepatitis in ferrets. J Virol 78, 12672–12676, doi:10.1128/JVI.78.22.12672-12676.2004 (2004).

9 Bolles, M. et al. A double-inactivated severe acute respiratory syndrome coronavirus vaccine provides incomplete protection in mice and induces increased eosinophilic proinflammatory pulmonary response upon challenge. J Virol 85, 12201–12215, doi:10.1128/JVI.06048-11 (2011).

10 Liu, L. et al. Anti-spike IgG causes severe acute lung injury by skewing macrophage responses during acute SARS-CoV infection. JCI Insight 4, doi:10.1172/jci.insight.123158 (2019).

## References

11 Dicks, M. D. et al. A novel chimpanzee adenovirus vector with low human seroprevalence: improved systems for vector derivation and comparative immunogenicity. PLoS One 7, e40385, doi:10.1371/journal.pone.0040385 (2012).

12 Bewig, B. & Schmidt, W. E. Accelerated titering of adenoviruses. Biotechniques 28, 870–873, doi:10.2144/00285bm08 (2000).

13 Maizel, J. V., Jr., White, D. O. & Scharff, M. D. The polypeptides of adenovirus. I. Evidence for multiple protein components in the virion and a comparison of types 2, 7A, and 12. Virology 36, 115–125, doi:10.1016/0042-6822(68)90121-9 (1968).

14 Corman, V. M. et al. Detection of 2019 novel coronavirus (2019-nCoV) by real-time RT-PCR. Euro Surveill 25, doi:10.2807/1560-7917.ES.2020.25.3.2000045 (2020).

15 Wolfel, R. et al. Virological assessment of hospitalized patients with COVID-2019. Nature, doi:10.1038/s41586-020-2196-x (2020).

